# Extending Bayesian Elo-rating to quantify the steepness of dominance hierarchies

**DOI:** 10.1101/2022.01.28.478016

**Authors:** Christof Neumann, Julia Fischer

## Abstract

The steepness of dominance hierarchies provides information about the degree of competition within animal social groups and is thus an important concept in socioecology. The currently most widely-used metrics to quantify steepness are based on David’s scores (DS) derived from dominance interaction networks. One serious drawback of these DS-based metrics is that they are biased, i.e., network density systematically decreases steepness values. Here, we provide a novel approach to estimate steepness based on Elo-ratings, implemented in a Bayesian framework (STEER: Steepness estimation with Elo-rating). Our new metric has two key advantages. First, STEER is unbiased, precise and more robust to data density than DS-based steepness. Second, it provides explicit probability distributions of the estimated steepness coefficient, which allows uncertainty assessment. In addition, it relies on the same underlying concept and is on the same scale as the original measure, and thus allows comparison to existing published results. We evaluate and validate performance of STEER by means of experimentation on empirical and artificial data sets and compare its performance to that of several other steepness estimators. Our results suggest that STEER provides a considerable improvement over existing methods to estimate dominance hierarchy steepness. We provide an R package `EloSteepness` to calculate the new steepness measure, and also show an example of using steepness in a comparative analysis.

## Introduction

Analyzing dominance relationships, dominance ranks and dominance hierarchies is a staple in studies of animal behavior. Results of such analyses feature prominently in the description of social structure in many animal species, spanning vertebrates as well as invertebrates.

One key aspect in this context is hierarchy steepness, which can be defined as “the size of the absolute differences between adjacently ranked individuals in their overall success in winning dominance encounters” (de Vries et al., 2006, p. 585). A hierarchy is considered *steep* if these differences are large, and a *shallow* hierarchy is one in which these differences between individuals are small. Hierarchy steepness is therefore often also referred to as *dominance gradient* (e.g., Barrett et al., 1999), and systems with steep hierarchies are often termed despotic while shallow systems are referred to as egalitarian, or less despotic (Sterck et al., 1997; Vehrencamp, 1983). Steepness is particularly relevant for questions related to socioecology, dominance styles, biological markets and phylogenetic covariation of social traits (Amici et al., 2020; Balasubramaniam et al., 2012a; Flack & de Waal, 2004; Schino & Aureli, 2008; Sterck et al., 1997; van Schaik, 1989).

For example, Balasubramaniam et al. (2012a) investigated variation in hierarchy steepness in groups of nine species of macaque (genus *Macaca*). They found evidence for a strong phylogenetic signal in hierarchy steepness, which is exceptional given the generally low magnitude of phylogenetic signals in behavioral traits (Blomberg et al., 2003; Kamilar & Cooper, 2013). Their results also fit a wider literature on phylogenetic covariation in a suite of behavioral traits in this genus (Balasubramaniam et al., 2012b; Matsumura, 1999; Thierry et al., 2000, 2008). This result helped to reconcile the influence of both, phylogenetic history and environmental factors, in shaping variation in behavioral patterns across species (Balasubramaniam et al., 2012a).

Quantifying hierarchy steepness has predominantly been done with one index (referred to here as ‘classic steepness,’ de Vries et al., 2006). This index, however, suffers from one important drawback. Specifically, the steepness index decreases as data density decreases, i.e., the more dyads that have no observed interactions in the dominance network, the shallower the hierarchy steepness (e.g., figure 2 in Klass & Cords, 2011) (see also figure A1). In other words, the higher the proportion of unknown relationships in a data set (the sparser the data set is), the smaller the steepness index becomes (see also Balasubramaniam et al., 2012a).

A new or improved index that does not suffer from this issue is therefore desirable. Any such index should have at least the following properties. First, it should indeed capture the phenomenon it is supposed to capture, i.e., the average difference in power differentials among individuals. Second, it should be robust to data density and observation effort. An additional desirable feature would be that the index also assesses the uncertainty arising from varying data density and observation effort. Here we propose a novel index to quantify hierarchy steepness that meets all three of these criteria. We begin by briefly describing the original steepness index of de Vries et al. (2006), followed by a description of our proposed novel index.

### Classic steepness based on David’s scores

Formally, steepness has been quantified as the slope of a regression of cardinal dominance scores of individuals on their ordinal dominance ranks, where the cardinal dominance scores are typically normalized David’s scores (de Vries et al., 2006). Calculating classic steepness (per de Vries et al., 2006) starts from a square matrix in which dyadic dominance interactions are tabulated. These raw interaction frequencies are then transformed into dyadic win/loss proportions (*P*_*ij*_). For example, if the dyad *AB* interacted 10 times, and *A* won 9 of these interactions, the winning proportion of *A* is *P*_*AB*_ = 0.9, and for *B* it is *P*_*BA*_ = 1 *− P*_*AB*_ = 0.1. Losing proportions are calculated analogously. These winning and losing proportions are then summed for each individual, being weighted by the winning and losing proportions of the opponents (see de Vries et al. (2006) for more details). The result of this procedure is a single score for each individual (David’s score, David, 1987; Gammell et al., 2003) where high values indicate high success (“high rank”) and low scores indicate low success (“low rank”). During the next step, David’s scores are then normalized such that the scores of all individuals range between 0 and *n −* 1, where *n* is group size. To derive the steepness metric, a simple linear regression is fit between scores and ordinal ranks of the scores (figure 1 in de Vries et al. (2006), see also figure 3). The absolute value of the slope coefficient of this model then is the steepness metric. Because of the normalization step, steepness ranges between 0 and 1.

**Figure 1:**
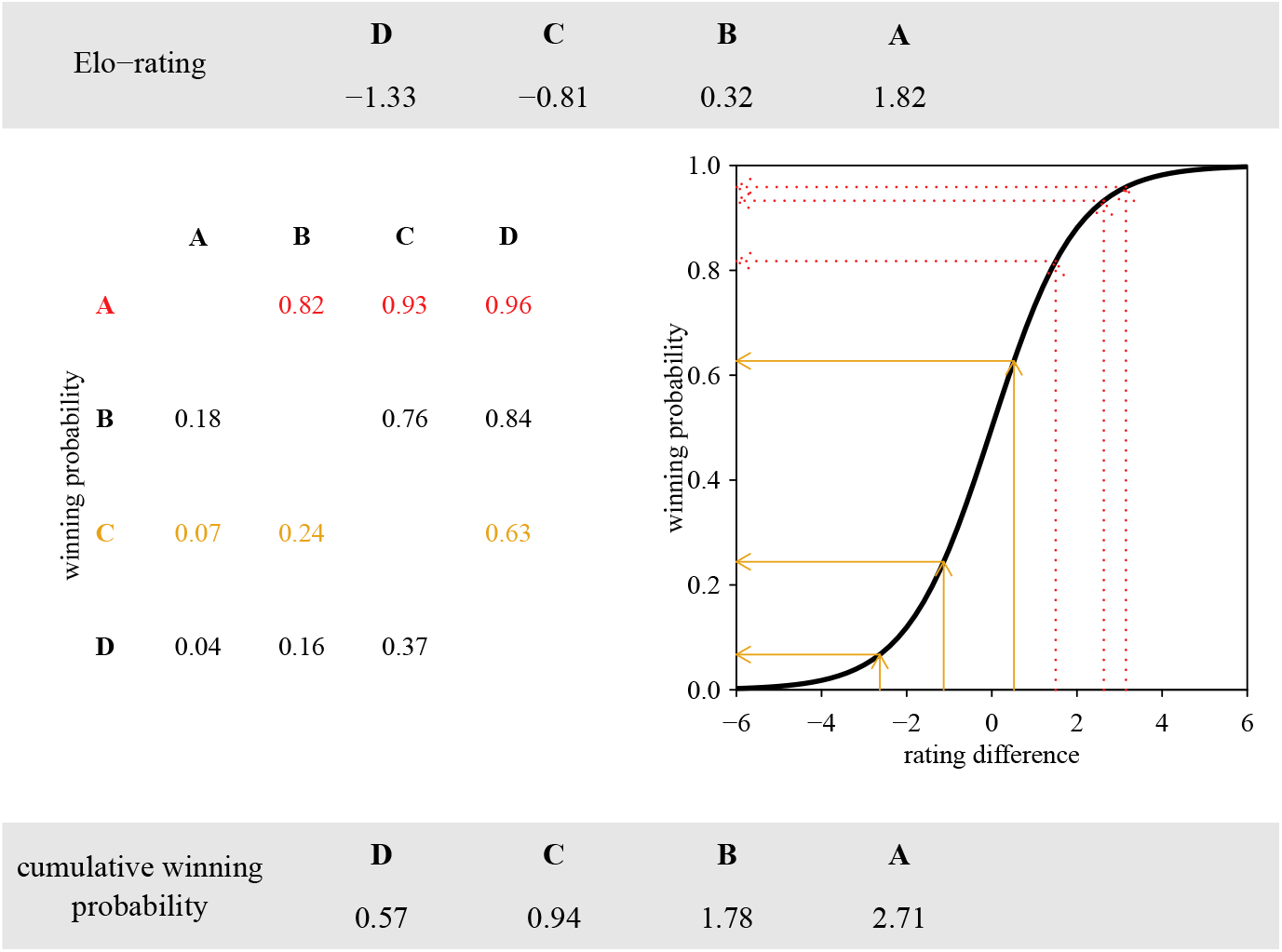
From individual ratings to cumulative winning probabilities. From individual ratings (top panel), rating differences are calculated. These differences translate (right plot) into dyadic winning probabilities, which can be tabulated (left matrix). Winning probabilities are then summed for each individual to derive cumulative winning probabilities (bottom panel). Two individuals are highlighted (A in dashed red, C in full gold). The winning probabilities of each individual against itself are omitted (see main text).

A variant of this approach also suggested by de Vries et al. (2006) uses dyadic proportions that are corrected for chance (*D*_*ij*_), rather than the pure dyadic winning proportions *P*_*ij*_ to derive David’s scores. Throughout the evaluation of our new method, we consider both variants of classic steepness.

### Elo-rating based steepness

Our approach to obtain a steepness index follows the same general logic: We use dyadic interactions to derive normalized individual dominance scores, which are used to fit a simple regression model, the slope of which then is the steepness metric. The key new feature is that the individual scores are derived from Elo-ratings rather than being based on David’s scores.

In brief, Elo-rating in its original form updates individual ratings after single observed dyadic interactions, where interaction winners increase their ratings and losers decrease their ratings (Albers & de Vries, 2001; Elo, 1978; Goffe et al., 2018; Neumann et al., 2011). The magnitude of change in the ratings depends on the expected outcome of the interaction, which in turn depends on the rating difference between the two interactants prior to the interaction. For example, if individuals *A* and *B* interact and prior to their interaction, *A* had a much larger rating than *B*, the expectation is that *A* is very likely to win the interaction, and conversely *B* is very unlikely to win. If *A* indeed wins the interaction then the updated ratings will change very little for *A* and *B*, if at all. If, contrary to the expectation, *B* wins the interaction, ratings will change substantially.

In addition to the expected outcome, the exact amount by which ratings change depends on the parameter *k*, which determines the maximum amount of change in ratings after a single interaction (Franz et al., 2015; Goffe et al., 2018). Typically, *k* is unknown and set to some arbitrary value like 100 (e.g., Neumann et al., 2011) (see also below).

Importantly, the Elo-rating algorithm treats interactions sequentially, i.e., individual ratings are updated after each interaction in the temporal order in which they occur/are observed. This constitutes a major difference to static methods like David’s score where the sequence of interactions is ignored. However, since we want to compare the performance of our Elo-rating-based method to that of static methods, we employ a randomization approach. For this, we translate static interaction matrices into randomized sequences in which the interactions may have occurred. This is necessary because the actual sequence of interactions is not available in static networks, which is the case when matrices are the data source (see also Sánchez-Tójar et al. (2018) and Clark et al. (2018)).

For our purposes, we take advantage of the fact that with Elo-rating, we can express the expected winning probability for any individual with any potential opponent at any point in the rating process, which is simply a function of the differences in ratings between the two individuals. Importantly, these expected winning probabilities are defined regardless of whether two individuals actually interacted.

It is also important to note that there is no consensus about the exact shape of the relationship between rating difference and winning probability (e.g., Franz et al., 2015; Goffe et al., 2018; Neumann et al., 2011; Sánchez-Tójar et al., 2018), although all implementations have in common a sigmoidal shape. Here we follow Goffe et al. (2018)^1^ and define the expected winning probability of *A* against *B* as

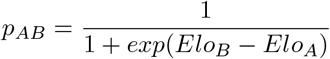

For example, consider the four individuals in figure 1. The ratings in the top row are the ratings after all interactions in the sequence have been evaluated. Here *A* has the highest rating among the four individuals and we can calculate *A*’s expected winning probabilities with the three remaining individuals. Since *A*’s rating is the highest rating of all individuals, all its winning probabilities are larger than 0.5 (dashed red arrows in figure 1), and in fact turn out to be all larger than 0.8. For example, for individuals *A* and *C*, with ratings of 1.82 and *−*0.81 respectively, we find that

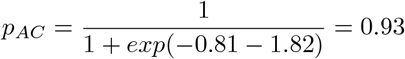

Individual *C*, on the other hand, is expected to win only against one other individual (*D*) and is expected to lose against *A* and *B*. This translates into one expected winning probability larger than 0.5 and two smaller than 0.5 (full golden arrows in figure 1).

These expected winning probabilities are then tabulated (center left panel in figure 1) per individual and opponent. When summing the winning probabilities per individual we obtain cumulative winning probabilities. These cumulative winning probabilities are akin to normalized David’s scores in that they range between 0 for an individual that has winning probabilities of 0 against all other individuals and *n −* 1 for an individual that is expected to win against all other individuals, where *n* is group size. More formally, we define individual *i*’s cumulative winning probability *c*_*i*_ as

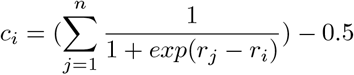

where *r*_*i*_ is individual *i*’s rating at the end of the interaction sequence, *r*_*j*_ is individual *j*’s rating at the end of the interaction sequence and *n* is the group size. We need to subtract 0.5 from this sum to account for the winning probability when *i* = *j*, i.e., the winning probability of individual *i* against itself, which is 1*/*(1 + *exp*(0)) = 0.5 and irrelevant.

With these cumulative winning probabilities at hand, we fit a regression model analogously to de Vries et al. (2006) with cumulative winning probabilities as a function of ordinal ranks of cumulative winning probabilities. The absolute value of the regression slope then is the steepness index (see also figure 3).

### Tackling uncertainty

So far, we defined Elo-based steepness as a point estimate. There are however several sources of uncertainty for obtaining this estimate. First, we usually do not know the ratings of all individuals at the start of the interaction sequence. Second, we do not know the value of *k*, which determines how much exactly ratings change after each interaction. Both these issues have been addressed before (Franz et al., 2015; Goffe et al., 2018; e.g., Newton-Fisher, 2017), and we follow Goffe et al. (2018), who adopted an explicitly Bayesian approach to estimate start ratings and *k* from the available interaction data, rather than setting them to arbitrary values. Consequently, all quantities of interest, such as start ratings and *k* can be seen as probability *distributions* rather than fixed point estimates. The same is true for to the expected dyadic winning probabilities and, in extension, to the cumulative winning probabilities of individuals.

One additional source of uncertainty concerns the actual sequence in which the interactions occurred. Recall that we translate static matrices, i.e., interactions that were aggregated over some time frame, into “dynamic” interaction sequences in order to apply Elo-rating (Clark et al., 2018; Sánchez-Tójar et al., 2018). As we often do not know the actual sequence, we simply use randomized versions of sequences in which the interactions could have occurred. The resulting multiple probability distributions can then simply be combined.

The concept is illustrated in figure 2. Beginning from an interaction matrix we obtain cumulative winning probabilities. In contrast to figure 1, these are now distributions rather than point estimates, stemming from MCMC samples. The final step in obtaining steepness is then to rank the cumulative winning probabilities and fit a simple linear regression. This procedure is applied in each of the MCMC samples separately and results in a probability distribution of steepness values.

**Figure 2:**
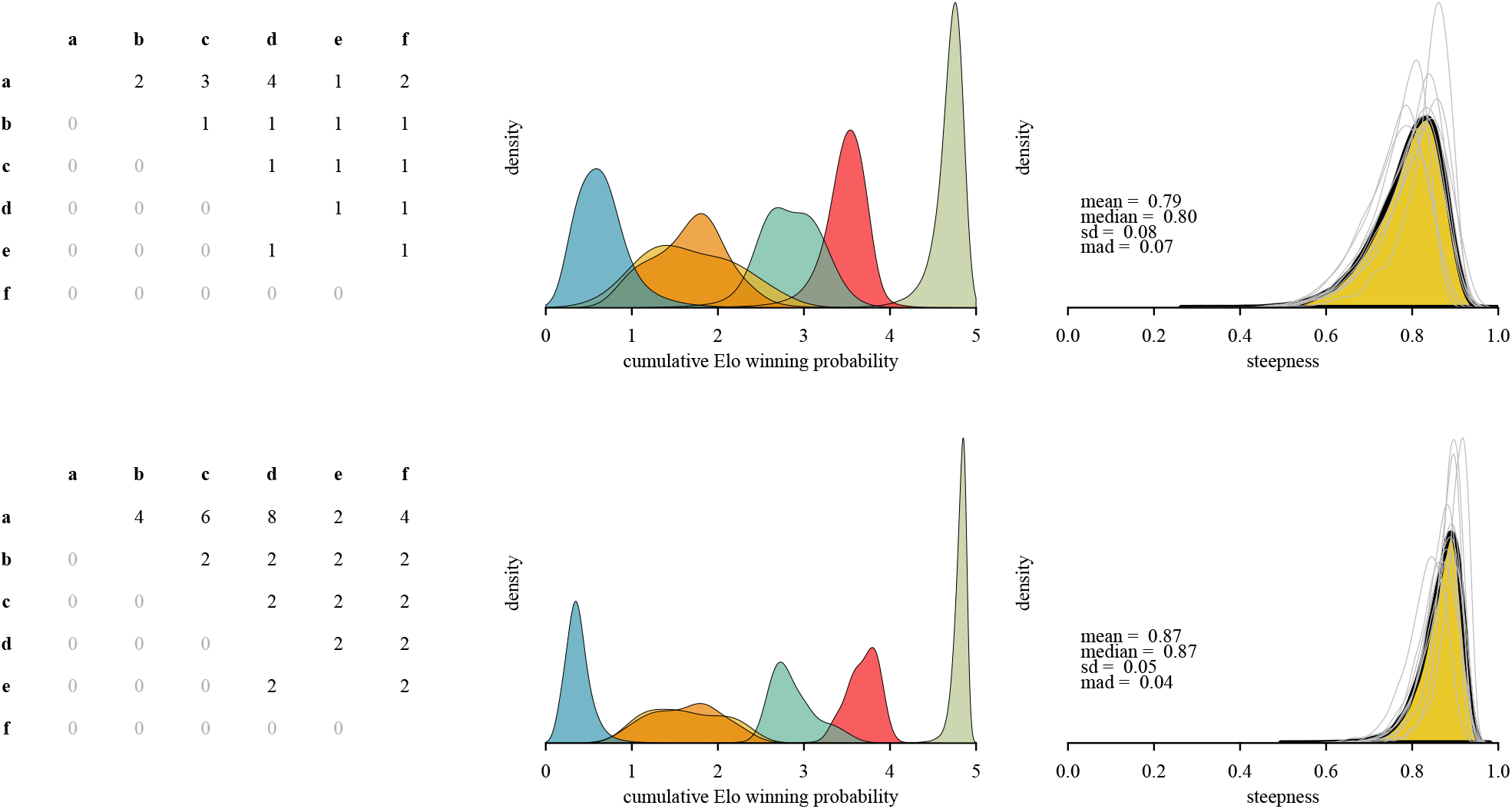
From interaction matrix to steepness distribution via cumulative winning probability distributions. The interactions were modeled with 10 randomized sequences and 4,000 MCMC samples each. In the center column the 40,000 samples are combined to display cumulative winning probabilities of individuals. In the steepness distribution in the right column, the 10 randomizations are depicted individually via the grey lines and combined via the yellow area. The lower panel replicates the upper panel with the difference that it is based on doubling the observed interactions.

Figure 2 shows that the amount of data directly informs how uncertain we need to be regarding the steepness estimates. Distributions shown in the bottom row vs. the top row of figure 2 have the exact same win/loss ratios, but differ with respect to the number of observed interactions: the bottom row has twice the number of observed interactions. With more observed interactions the resulting distributions become narrower, as they should. The more information (here: observed interactions) we have, the less uncertain we can be about our estimates. Another way of looking at this is that the cumulative winning probability distributions in the lower panel overlap much less compared to the upper panel.

It is also noteworthy that using the Bayesian Elo-model in this context handles ambiguous cases very naturally. In the example here, individuals *d* and *e* have a tied relationship, i.e., they both won and lost the same number of interactions with each other. As a result, their cumulative winning probability distributions overlap to a very large extent.^2^

In summary, our new steepness measure follows an analogous approach as classic steepness by using standardized scores of individuals as the source to estimate the steepness slope (figure 3). In contrast to classic steepness though, these individual scores represent probability distributions of cumulative winning probabilities derived from Elo-ratings. We therefore refer to it as STEER: steepness estimation with Elo-rating. STEER captures uncertainty on multiple levels: first, the uncertainty arising from the sequence itself, i.e., by randomizing the order in which interactions are considered; second, from the actual rating process, i.e., by using Bayesian estimates of *k* and start ratings (Goffe et al., 2018).

**Figure 3:**
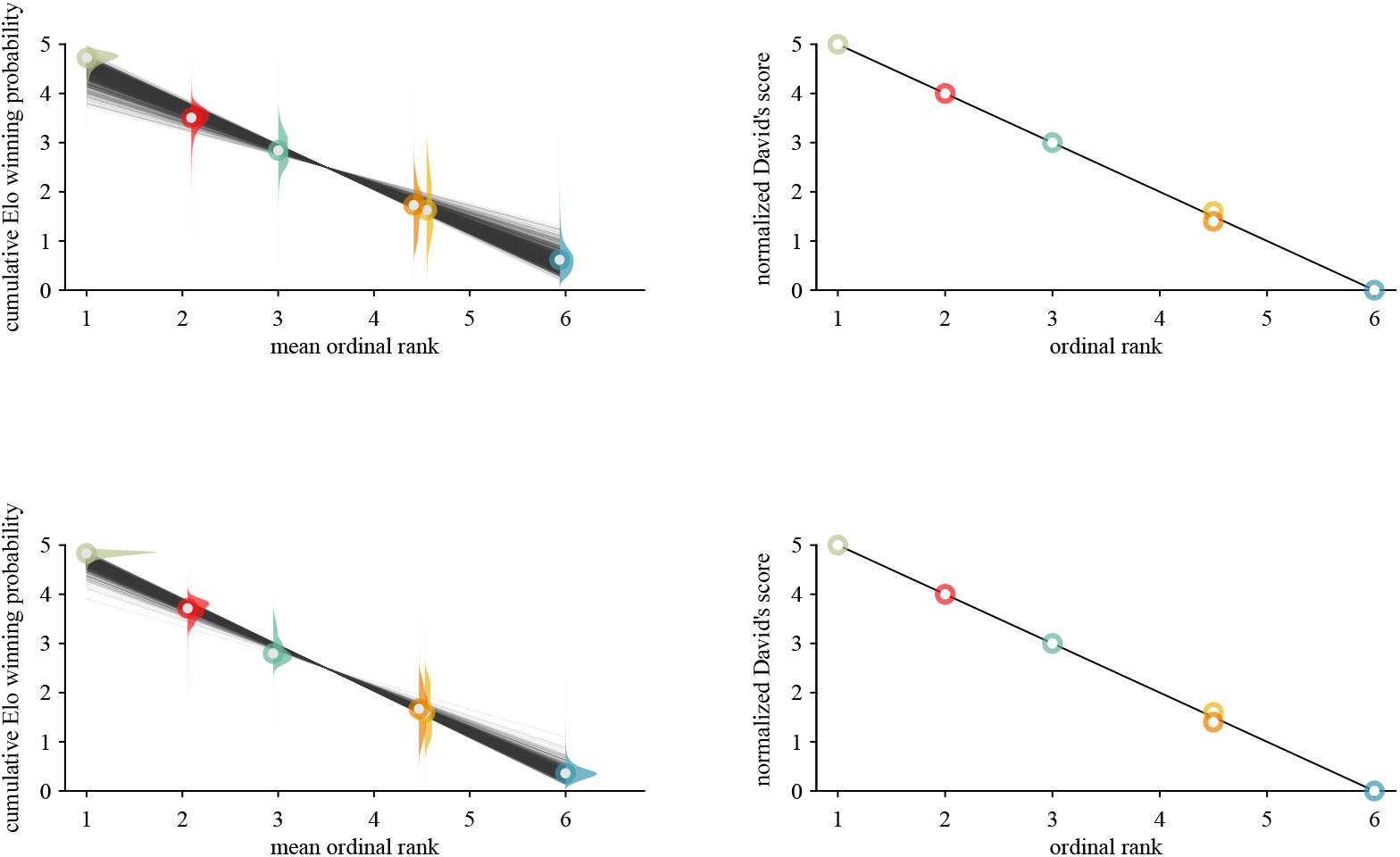
Elo-based steepness and classic steepness. Classic steepness (right column) is calculated with *P*_*ij*_ David’s scores and represents a point estimate. Two individuals have the same David’s score and hence have a tied rank and therefore are slightly jittered vertically for visual purposes. Elo-based steepness carries over uncertainty deriving from, amongst others, data density. The lower panel has twice the number of interactions observed compared to the top panel, which informs the Bayesian Elo steepness, but has no effect on classic David’s score steepness.

## Material and methods

We used two data sets, which we describe in more detail below. The first was a set of artificially generated matrices. The second was a set of empirical data sets, i.e., matrices extracted from published sources.

With these data sets, we used two complementary approaches to evaluate the performance of STEER in comparison to the original (classic) steepness measure and three other options to quantify steepness.

First, we looked at reliability by assessing how well the different methods recovered some underlying (‘true’) steepness value. We used two ways of defining ground truth. For the artificial data, we knew ground truth because we set the steepness parameter during data generation (see below and figure A2). For the empirical data, this approach was not possible and we resorted to using steepness derived from *P*_*ij*_ David’s scores as ground truth. Given the known detrimental effects of unknown relationships on this steepness measure (Klass & Cords, 2011), we only used dense interaction networks (less than 5% unknown relationships) for this analysis. For comparison, we also used this second approach (considering steepness from *P*_*ij*_ David’s scores as ground truth) to evaluate method performance for the artificial data even though we knew the actual ground truth. An ideal method would have had a correlation *ρ* = 1 (‘precise’) and a regression slope *β* = 1 (‘unbiased’) when looking at the relationship between ground truth and the results of the different methods to assess steepness.

Second, we investigated how well the different methods dealt with sparse data. Here, we performed a removal experiment, in which we increased the proportion of unknown relationships incrementally in a given data set by removing interactions, and subsequently quantified the relationship between steepness and the proportion of unknown relationships in each data set (see figure A3). This approach provided insights into the robustness of the different methods with respect to data density. Specifically, we started by assessing the proportion of unknown relationships in the initial network. Then we removed interactions from the network until one more dyad had no interactions, which corresponds to an increase in the proportion of unknown relationships, and stored the resulting matrix. Then we repeated the removal procedure until we reached a matrix with 70% unknown dyads (which represents a fairly empty/sparse network) and stored each intermediate matrix. For example, a matrix with 10 individuals and hence 45 dyads, with all dyads initially observed, could have 31 dyads removed until reaching 70% unknown dyads. This would result in 32 matrices (31 removal steps + the initial matrix). In order to speed up computation, whenever the number of resulting matrices was larger than 12, we randomly selected 12 matrices and discarded the remaining ones.

The quantity of interest in this experiment was the relationship between steepness and the proportion of unknown relationships. For doing this, we separately fitted a linear regression for each data set (set of up to 12 matrices) and for each steepness algorithm. The steepness value was the response variable and the proportion of unknown relationships was the predictor variable. The slope of this regression provided information on how a method responds to unknown relationships.

A perfect method would produce the same steepness measure regardless of the proportion of unknown relationships, and hence the slope would be zero, i.e., a method returns (on average) the same steepness for each value of unknown relationships. If the slope is positive, the steepness value becomes larger with increasing unknown relationships, i.e., sparser matrices. If the slope is negative, the steepness value becomes smaller with increasing unknown relationships. This latter pattern is what we expected for the classic steepness metrics, and we expected it to be more pronounced, i.e., more negative, in the *P*_*ij*_-based steepness compared to the *D*_*ij*_-based steepness (de Vries et al., 2006).

### Data sets

#### Artificial data sets

We generated 1,000 artificial dominance interaction matrices. First, we set a group size between 5 and 25, which corresponds to the range of the majority of published dominance interaction matrices (Neumann, 2022b). Then we set the number of interactions dependent on group size with 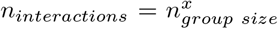, where *x* was a random number from a uniform distribution ranging between 1.8 and 2.8. For a matrix with five individuals, this lead to interaction numbers ranging between 19 and 91, and for the largest groups with 25 individuals to between 329 and 8208 interactions. The starting steepness for each matrix was set to a random, uniformly distributed, number between 0.2 and 1 (figure A2).

We also introduced two kinds of biases in how interactions were distributed across dyads. First, interaction frequency depended on how close two individuals were in rank. This parameter ranged from all dyads having the same underlying propensity to interact regardless of rank distance to situations where individuals of adjacent ranks interacted more frequently than pairs of individuals with large rank distances (e.g., Jennings et al., 2006; Seyfarth, 1976). The second bias allowed individuals interacting from equally often to some individuals interacting more often than others.

#### Empirical data sets

We compiled a database that contained 978 published dominance interaction matrices. Of these, we only kept those that met the following criteria: The number of individuals was at least five, the proportion of unknown relationships was less than 0.5 and all individuals were observed in at least one interaction. After this selection process 670 unique matrices remained (Neumann, 2022b).

### Algorithms used

In addition to our new steepness measure, we also subjected five other algorithms/variants to our evaluations. The first were the two versions of classic steepness, based on *D*_*ij*_ and *P*_*ij*_ winning proportions as described above (de Vries et al., 2006). We refer to these two as 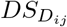 and 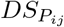.

The third steepness metric was based on a Bayesian version of David’s scores, which we also developed in the course of this study and describe in more detail in the appendix. We refer to this method as *DS*_Bayes_.

The fourth algorithm was based on repeatability of Elo-ratings (Sánchez-Tójar et al., 2018). Here an interaction matrix is translated into a large number, typically 1000, of randomized interaction sequences. Each sequence is then subjected to the Elo-rating algorithm, which results in a set of multiple ratings for each individual (one rating per individual per randomized sequence). For these ratings, repeatability (also known as the intra-class correlation coefficient, Nakagawa & Schielzeth, 2010) is calculated, which serves as steepness estimate (Sánchez-Tójar et al., 2018). We refer to this method as Elo_rpt_.

The final algorithm was simply the proportion of interactions that go against the rank order (*upward steepness*). To this end, an ordinal ranking of individuals is produced first. Here we use classic David’s score to produce this ranking. Then the interaction matrix is reordered according to the obtained ranking. The upward steepness is just the proportion of interactions below the diagonal divided by the total number of interactions. It is noteworthy to say that this index can, at least theoretically, be zero if the produced ranking is completely false, i.e., the initially produced ranking is the opposite of the ‘true’ ranking and there are no entries above the diagonal in the dominance matrix.

All algorithms share the same scale, i.e., they all range between 0 and 1 where 0 indicates a shallow hierarchy and 1 indicates a maximally steep hierarchy. This feature makes the comparison between the different methods straight-forward. Also note that although STEER and steepness based on Bayesian David’s scores produce posterior *distributions*, we reverted to using posterior medians (i.e., point estimates) in the quantitative evaluations to simplify the comparison.

### Software

All data were generated and analyzed with the `EloSteepness` (Neumann, 2022a) and `EloRating` (Neumann & Kulik, 2020) packages, which are based on `Stan` (accessed through `rstan` and `cmdstanr` (Stan Development Team, 2020)) and `Rcpp` (Eddelbuettel & Francois, 2011). Steepness based on repeatability was obtained from the `aniDom` package (Farine & Sánchez-Tójar, 2021). All data we used and generated are in the `EloSteepness.data` data package (Neumann, 2022b).

## Results

### Recovering ground truth

All methods showed a fairly tight and, with the exception of Elo_rpt_, linear relationship between the estimated steepness and the steepness we set during the data generation (figure 4). Upward steepness consistently produced higher steepness values than expected, although the relationship was still a linear one.

**Figure 4:**
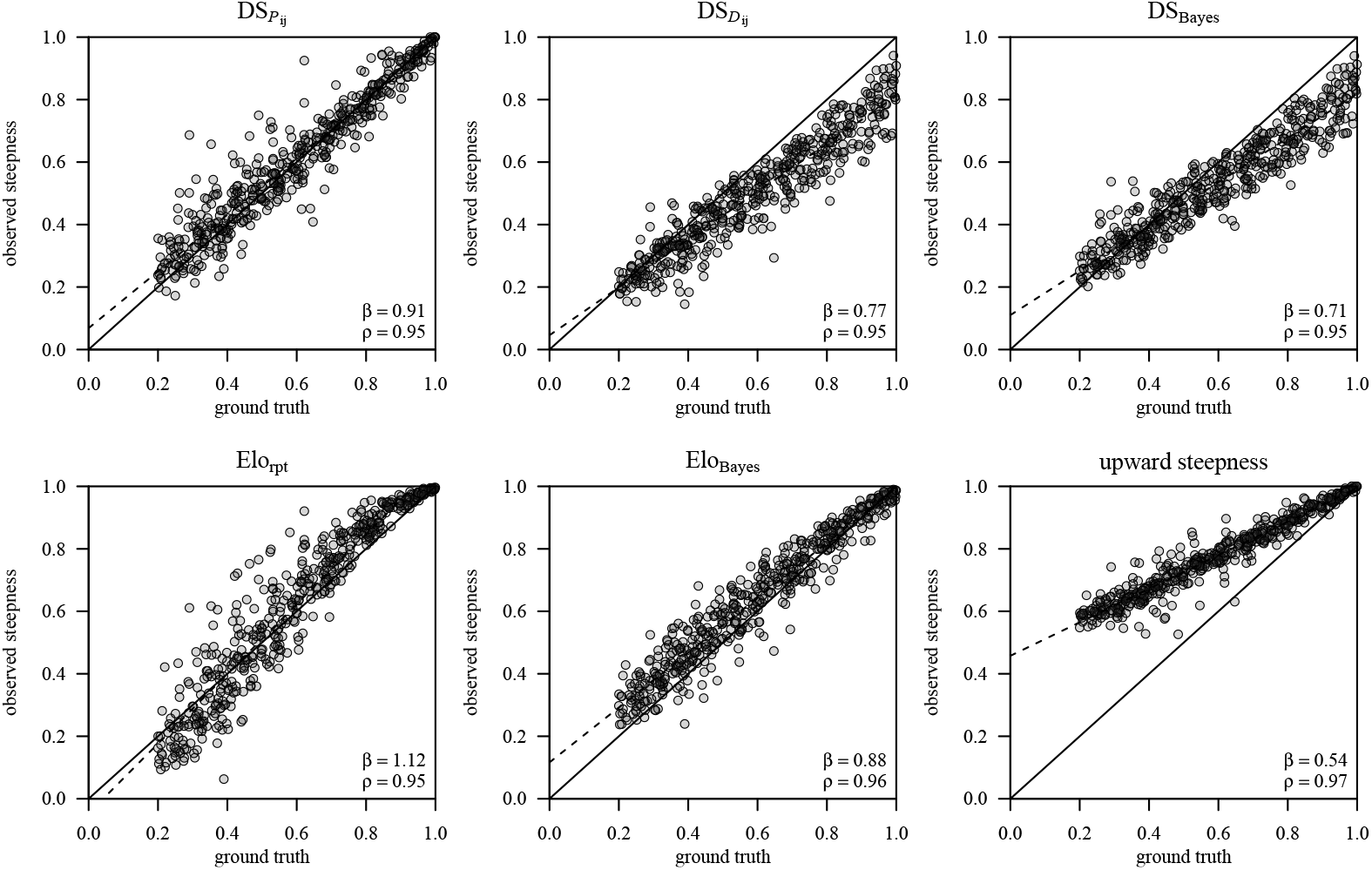
Relationship between estimated steepness and ground truth for 1000 `artificial` data sets. *Ground truth* here refers to steepness as we set it during the generation of the artificial data. Only data sets with less than 5% of unknown relationships are included. *β* is the slope estimate from a regression accross all data sets and *ρ* is the correlation coefficient (both should be ideally 1).

For the empirical data sets we did not know the ground truth and we resorted to using *P*_*ij*_ David’s score steepness as ground truth instead (figures 5 and 6). To be conservative, we used only data sets that had less than 5% unknown relationships, and for comparison we also applied this analysis to the artificial data. Since ground truth is unknown for the empirical data sets, the results presented in figures 5 and 6 have to be taken with some caution, because of the problems associated with the classic steepness measures pointed out above.

**Figure 5:**
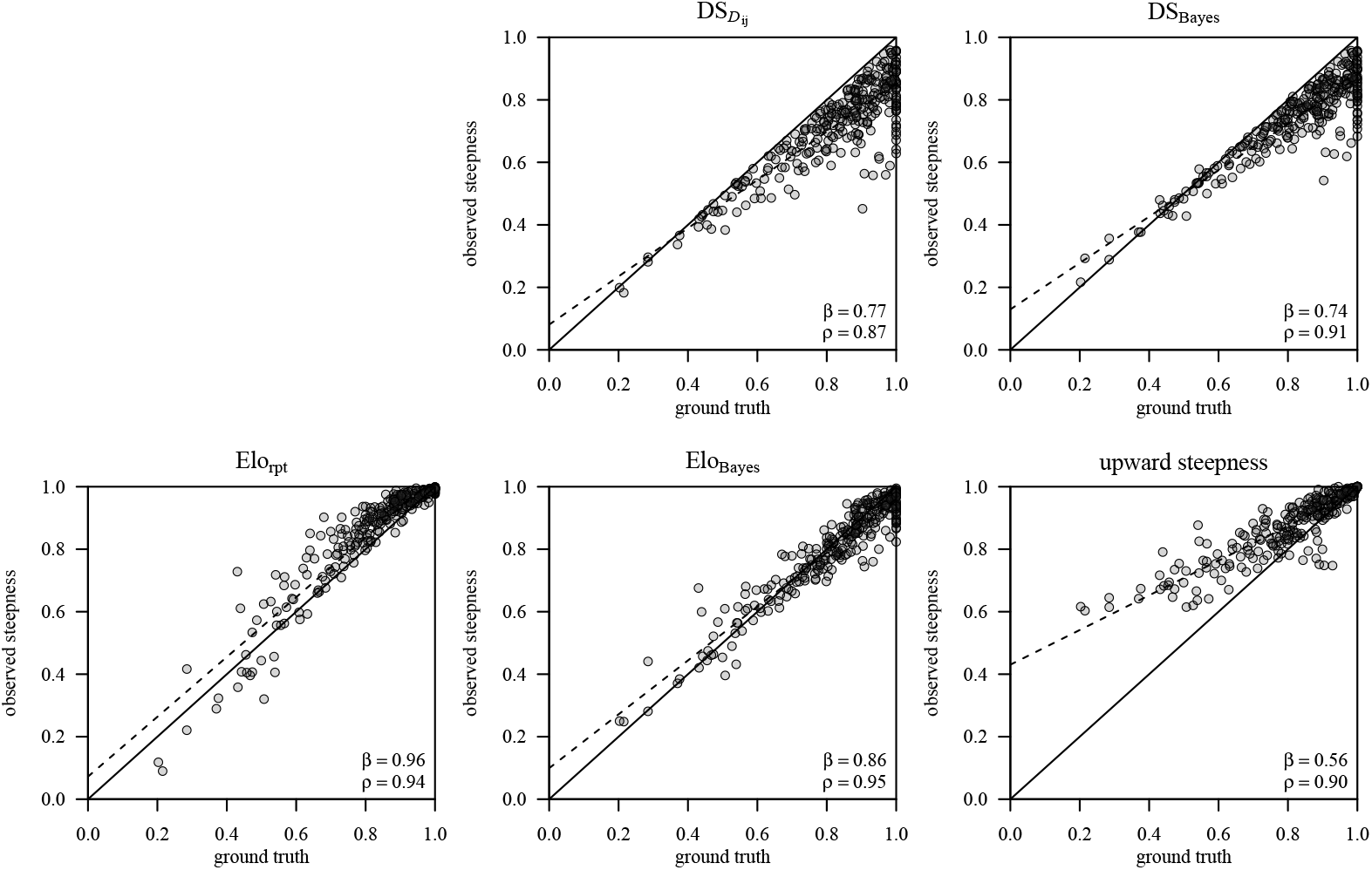
Relationship between estimated steepness and ground truth for 670 `empirical` data sets. *Ground truth* here refers to steepness as estimated with *P*_*ij*_ David’s scores. Only data sets with less than 5% of unknown relationships are included.

**Figure 6:**
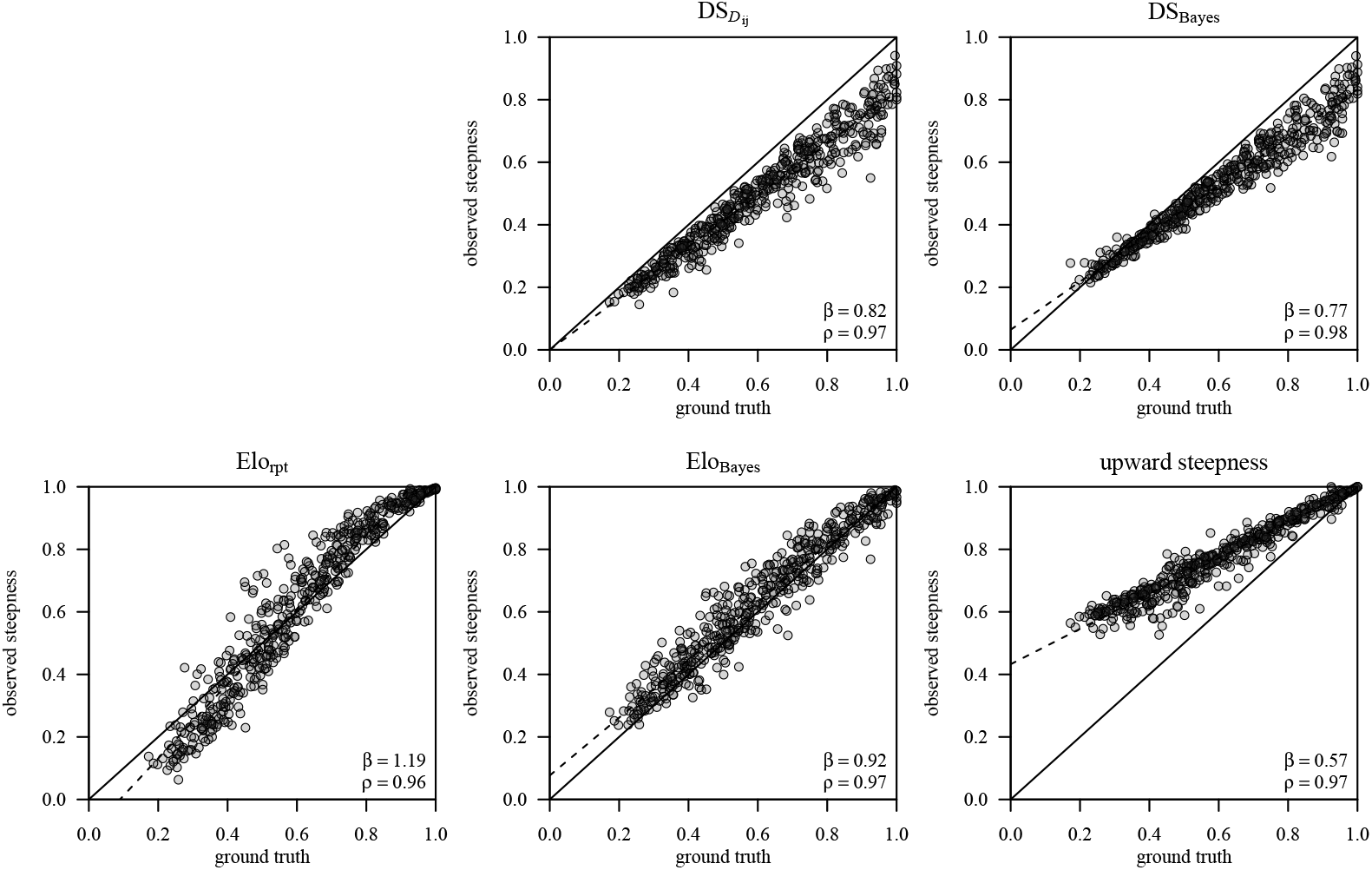
Relationship between estimated steepness and ground truth for 1000 `artificial` data sets. *Ground truth* here refers to steepness as estimated with *P*_*ij*_ David’s scores. Only data sets with less than 5% of unknown relationships are included.

For both the empirical and the artificial data, our new method produced on average the most accurate and least biased results, i.e., the recovered steepness showed a tight linear relationship with the ground truth. The remaining methods showed much stronger biases, either underestimating (classic steepness and Bayesian DS steepness) or overestimating (upward steepness), and in the case of Elo_rpt_ also showing a non-linear relationship with the ground truth.

#### Handling sparse data sets/removal experiment

All steepness methods based on David’s scores, including Bayesian David’s scores, showed a strong negative dependence (as expected) on the proportion of unknown relationships (figures 7 and 8), i.e., steepness decreased the sparser a data set became, which resulted in negative slopes (as depicted in the figures). In contrast, the two Elo-rating-based methods (including STEER) and the simple upward steepness showed on average little dependence on unknown relationships.

**Figure 7:**
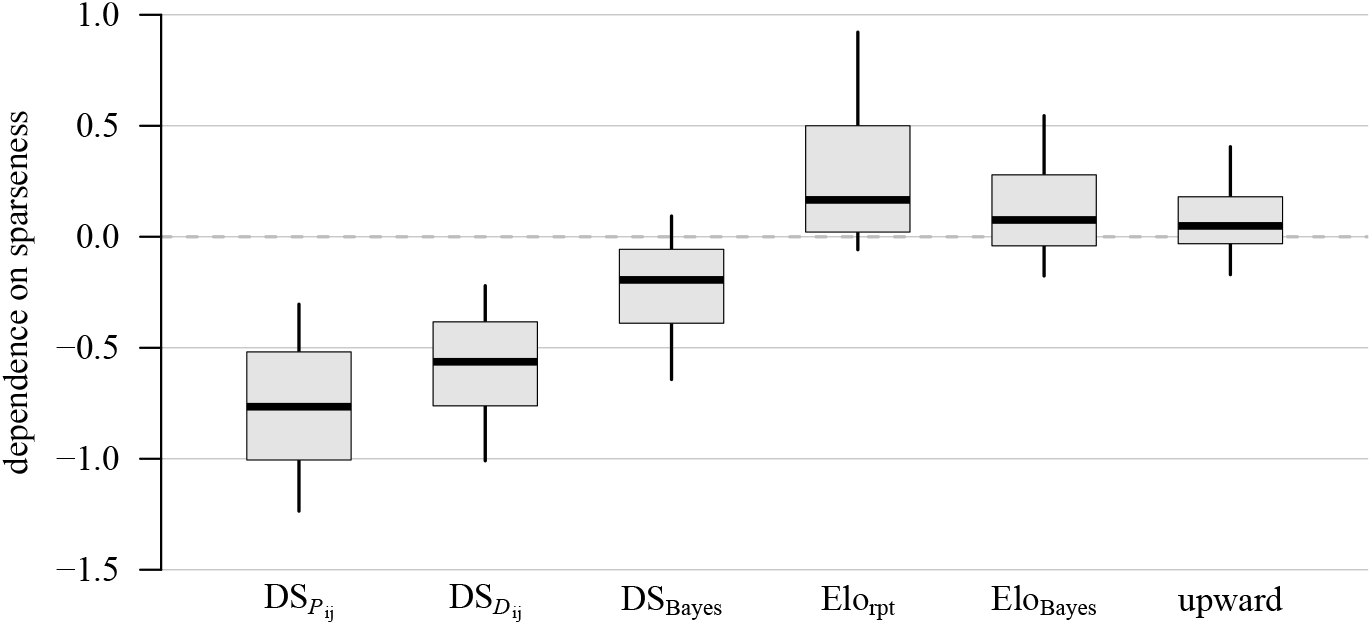
Dependence of steepness on unknown relationships with `artificial data`. A perfect method would have no dependence, i.e., slope of 0. All *n* = 1000.

**Figure 8:**
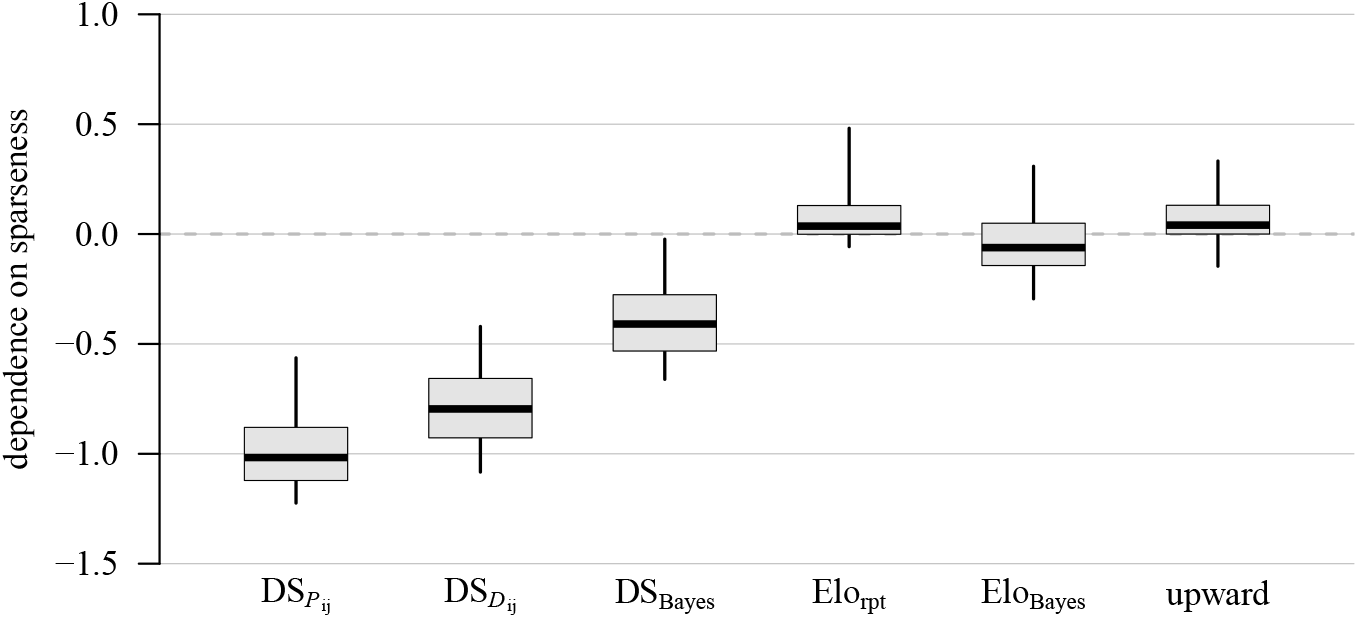
Dependence of steepness on unknown relationships with `artificial data`. A perfect method would have no dependence, i.e., slope of 0. All *n* = 670.

### An applied example

To illustrate a potential application of our new measure we performed an analysis similar to that of Balasubramaniam et al. (2012a) who studied variation in dominance hierarchy steepness in social groups of macaques (*Macaca* sp.). One of their main conclusions was that there is more variation in steepness between species than within species, which they interpreted as providing support for an substantial phylogenetic component in this particular facet of social behavior.

Here we ran a similar analysis, which nevertheless differed in several ways from that of Balasubramaniam et al. (2012a). First, we used a larger data set, which comprised more groups and species, and we also did not restrict the data to female-female interactions. Second, we estimated between-species variation and within-species variation in a single model, as opposed to running two separate sets of analyses. The response variable in our model was the STEER steepness estimate, which we considered to be beta-distributed. We estimated the two parameters of the beta distribution (mu and phi), and more importantly, two variance components. The first variance component reflected the phylogenetic relationships between species assuming a Brownian motion model, i.e., between-species variation, using a pruned tree from a consensus phylogeny of primates (Arnold et al., 2010). The second component reflects the variance due to repeated measurements of the same species, i.e., within-species variation. This model resembles a phylogenetic generalized linear mixed model (de Villemereuil & Nakagawa, 2014; Hadfield & Nakagawa, 2010). Note that the response variable itself actually represented posterior distributions of the steepness estimate as we included the estimation of STEER in the same model. In other words, we modeled steepness from the interaction data simultaneously with the within- and between-species variation in the same single model. We coded this model in Stan (Stan Development Team, 2020) (see Neumann (2022b) for data and model code).

We found no evidence for larger between-species variation (mean SD = 0.21) compared to within-species variation (mean SD = 0.20) (figure 9). Rather the posterior distributions of the two variance components overlapped substantially, therefore suggesting similarly sized magnitudes of within- and between-species variation.

**Figure 9:**
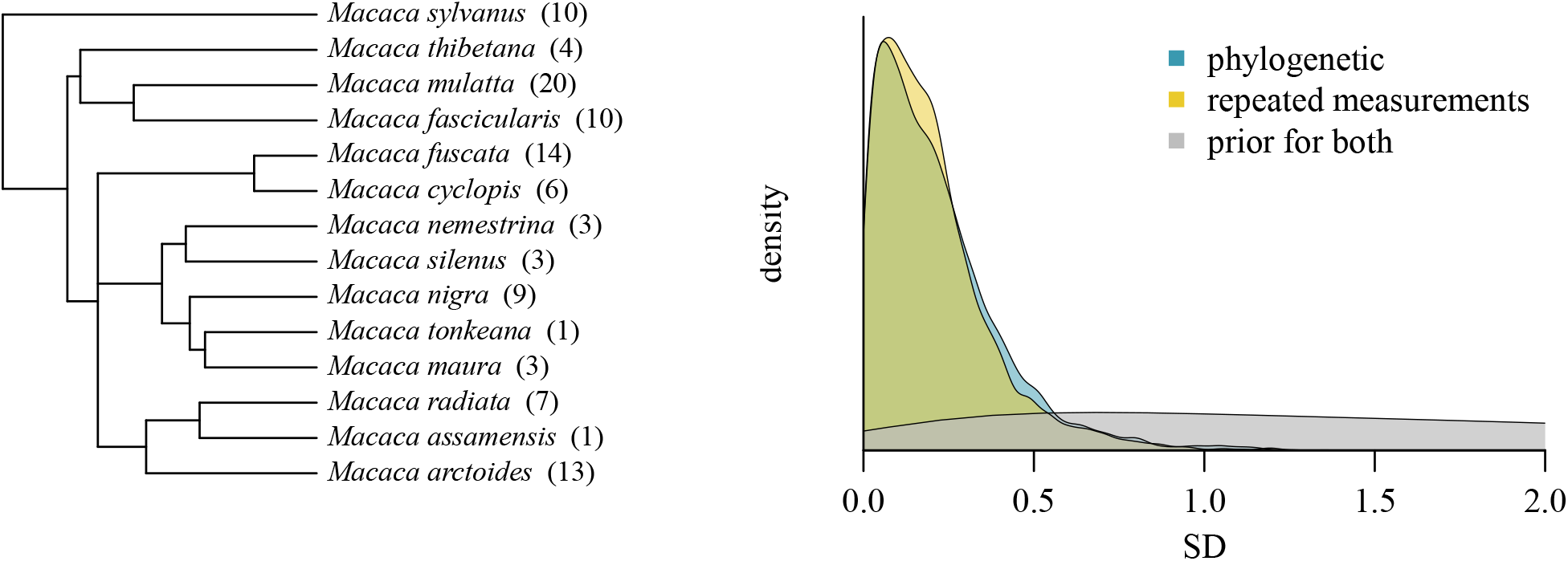
An example of applying STEER in a comparative study. Here we estimated steepness in 104 groups of 14 macaque species and assessed variance due to phylogeny and due to repeated measurements of groups of the same species. The two posteriors overlapped almost completely and differ only marginally in their central tendencies.

Note that these results are not to be taken as refuting the results and conclusions of Balasubramaniam et al. (2012a) because there are several important methodological differences between our and their analyses (for example we did not constrain our data to using exclusively data on adult females) (see also figure A4). Rather, the point of this example is to illustrate the potential of our new method to answer these kinds of questions.

## Discussion

In this study we presented STEER, a novel algorithm to quantify steepness of animal dominance hierarchies and evaluated its performance in comparison to other available methods. The new method based on Bayesian estimation of Elo-ratings is a considerable improvement over existing algorithms to estimate hierarchy steepness. It recovered ground truth faithfully and it showed little systematic dependence on unknown relationships. Among the methods we tested, it was the only one that performed well in both these contexts, i.e., being precise and unbiased. In contrast, the second method we developed, based on a Bayesian implementation of David’s scores still outperformed classic steepness with respect to precision and bias, but performed overall less well than STEER.

It is noteworthy that for our evaluation we ignored the fact that our method’s output represents a Bayesian posterior distribution of steepness. Rather, we treated it here as a point estimate (the median of the posterior distribution) to simplify the comparison with the other methods, which provide point estimates. However, it is clearly beneficial to have a sense of uncertainty of steepness, which our new method provides via credible intervals, which makes inference about uncertainty much more explicit and straightforward. For example, a 89% credible interval for a median steepness of 0.8 that ranges from 0.31 to 0.97 should be taken more cautiously than an interval of 0.73 to 0.85 for the same median steepness. These uncertainties then can either be used descriptively when characterizing the hierarchy of an animal group, or can be carried forward in cases where steepness is a predictor or response variable in analyses that comprise multiple groups within or between species (e.g., Balasubramaniam et al., 2012a; Kaburu & Newton-Fisher, 2015; Schino & Aureli, 2008). We did the latter in our reanalysis of Balasubramaniam et al. (2012a) (figure 9, see also figure A4 for an analysis that uses point estimates of steepness).

Importantly, our method is currently implemented only for ‘static’ interaction networks, i.e., data sets for which the sequence of interactions is not known. However, it seems clearly beneficial to extend the approach to dynamic networks, where the sequence of interactions is actually known. Then it would be possible to account for (temporal) changes in individuals’ inherent fighting ability (‘rank changes’), or even changes at the group level. For example, it could be hypothesized that steepness should be larger when competition for resources becomes more intense. When this competition is measured quantitatively (e.g., number of receptive/fertile females available to males who compete for conceptions, availability of high-quality food or territories), then a temporal implementation would allow monitoring the relationship between steepness and competition without the need of dividing data sets into arbitrary time blocks.

### Performance of other methods

The two steepness measures based on classic David’s scores performed poorly in our evaluations. This result is not surprising because their biases and susceptibility to underestimating steepness are well-known (e.g., Klass & Cords, 2011) and provided the original impetus for the development of our new method. We therefore think it justified to suggest abandoning their use given that we now have more robust alternatives.

The steepness estimate provided by assessing repeatability of Elo-ratings has two downsides. First, it results in slightly biased steepness estimates by overestimating large and underestimating low steepness values. Second, there is no clear theoretical rationale for why repeatability of individual dominance scores should mechanistically provide us with an index that describes average differences in individual dominance success, i.e., steepness. While both arguments by themselves do not appear to be major issues, the combination of the two together with the availability of our new method also let us suggest to not use repeatability for steepness assessment.

The upward index, despite its simplicity, performed surprisingly well overall, although it tended to overestimate steepness. It must be noted though that it relies on ranking the individuals in the first place, which in itself is susceptible to biases. As such, its theoretical minimum of 0 can only be reached if the ranking that is initially required is completely wrong, which seems an unlikely outcome of any ranking algorithm known to us (e.g., Bayly et al., 2006; Neumann et al., 2018). As a result, it appears that the upward index is likely to overestimate steepness, which indeed seems to be the case (figure 4). Furthermore, one other potential drawback of this method is that it pools all interactions across dyads and hence might be susceptible to dyads that interacted disproportionately frequently.

All these options share the absence of a direct assessment of uncertainty because they are point estimates. As noted above, the steepness derived from Bayesian Elo-rating provides such an assessment in a very explicit fashion, which in itself is a major advantage.

Lastly, we also want to point to our implementation of Bayesian David’s scores. While its performance in the steepness context is not quite up to our new measure based on Elo-rating, it still performs better than the classic David’s scores. Given the popularity of DS to quantify individual dominance strength (in addition to forming the basis of classic steepness), it might be a fruitful follow up to properly validate these scores as a dominance measure in their own right.

### Guidelines/recommendations

When applying our new steepness measure we recommend to present the results in a way that does justice to its probabilistic (Bayesian) nature, i.e., provide readers with an appropriate assessment of the uncertainty associated with the outcome of the analyses. We thus recommend to provide numerical (mean, median, credible interval) as well as a graphical presentation of the posterior distribution of the steepness index.

Sánchez-Tójar et al. (2018) provided some guidelines regarding the required amount of data, i.e., the average number of recorded interactions per individual, that is necessary to infer a reliable dominance hierarchy. Specifically, they suggested that at least ten (better 20) interactions per individual need to be observed (Sánchez-Tójar et al., 2018, p. 605). We find it hard to make an analogous statement with regards to our new steepness measure. We want to avoid as much as possible recommending arbitrary cut-offs, which potentially force researchers to make binary decisions. Arguably, this could lead to under-reporting of data and results simply because a researcher who failed to reach a specific cut-off might choose to not report their results. Rather, we would advise to let the data ‘speak for themselves.’ The reason for this recommendation is that we conceive of steepness as a distribution rather than a point estimate, which allows assessment (via the width of the steepness distribution) of how confident we can be in our results given our observations. Seen like this, we should expect that more observed interactions lead to narrower steepness distributions. At the same time, we should *not* expect that more observed interactions lead to changes in the central tendencies (median or mean) of the steepness distribution.

Figure 10 shows these two relationships between amount of data (number of interactions divided by number of individuals) and point estimate (median of distribution), and between amount of data and the width of the 89% credible interval around the point estimate (as a direct measure of uncertainty of the point estimate). This figure illustrates that the point estimate is by and large independent of the amount of data, although it appears that very large steepness values are not likely to appear with low numbers of observed interactions. More importantly (and not surprisingly), the uncertainty decreases with increasing observations, i.e., credible intervals become narrower with larger numbers of observed interactions. Neither of these two plots suggests any clear cut-off. In other words, rather than using some more or less arbitrary cut-off to decide whether the observed steepness is an accurate description of the world, we find it more intuitive to let the data speak for themselves: a wide credible interval should make us less confident than a narrow credible interval.

**Figure 10:**
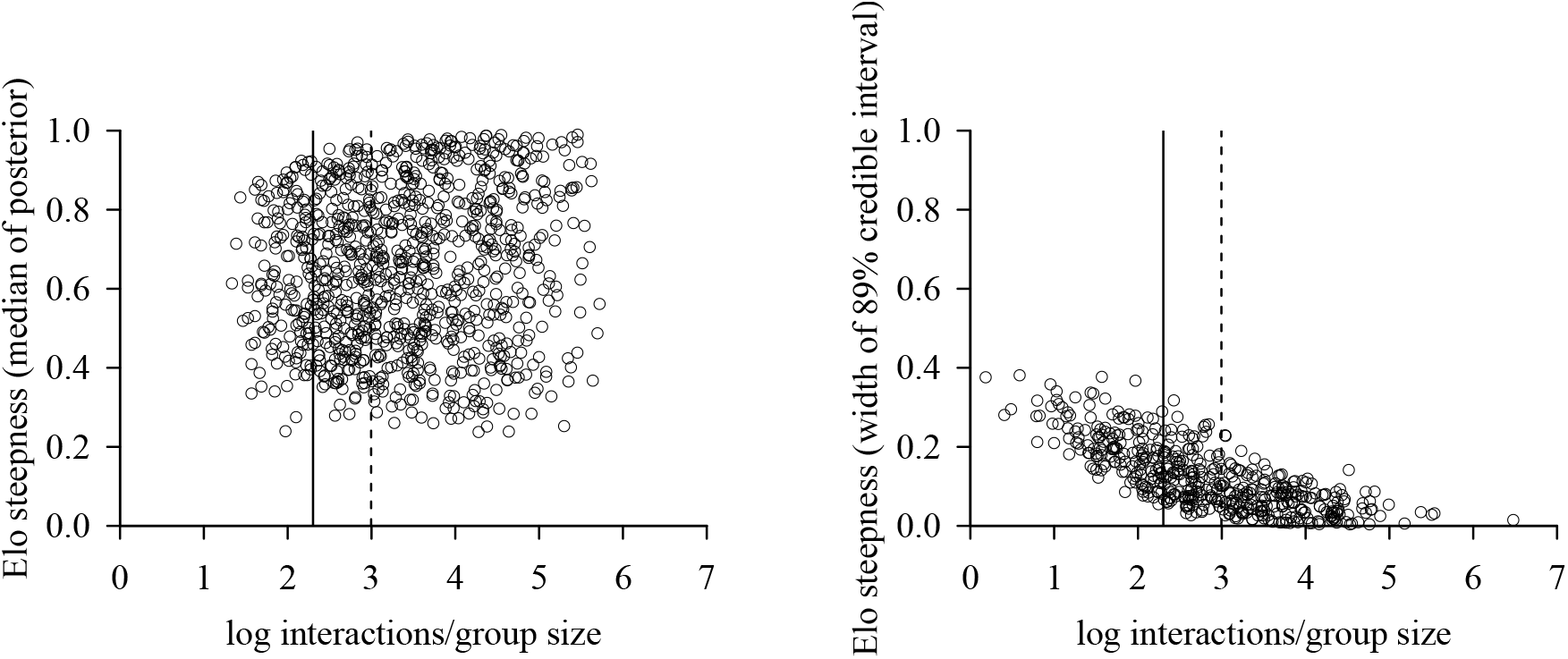
Point estimates and width of credible intervals of steepness as a function of number of observed interactions for artificial data sets. For reference, we included the recommendations put forward by Sánchez-Tójar et al.(2018) at ratios of 10 (full line) and 20 (dashed line) interactions per individual.

We do, however, agree with Sánchez-Tójar et al. (2018) in stressing the importance of *reporting* data characteristics of the studied networks (average number of observations per individual, proportion of unknown relationships). Ideally, this reporting is accompanied by the raw interaction data from which steepness was estimated.

The final issue that needs consideration is the number of randomized sequences to use when applying our new method. In principle, a simple rule would state that one would use as many randomizations as possible. For example, using 1,000 randomizations is a widely-used number in many contexts. For our evaluations this presented a practical problem, due to limited computational resources. We therefore established a rough guide, which uses 50 randomizations for matrices with less than 100 interactions, 20 randomizations for matrices with between 100 and 500 interactions, and 5 for matrices with more than 500 interactions. To establish this guide we took the following approach. For each data set, we randomly selected one matrix (which is either the full matrix, or one of the matrices from which interactions were removed). We then applied our algorithm 1,000 times to each matrix. As our ‘true’ steepness value we took the median of all posterior samples from 500 of those randomizations. From the remaining sequences we calculated steepness multiple times from one sequence, two sequences, five, ten, 20, 50, 100 and 200 sequences. We then checked at which of these increments the derived steepness differed less than 0.01 from our ‘true’ value. The result of this procedure gives an approximate and rough idea at which number the estimated steepness distributions levels off, i.e., at which value of randomizations do we get sufficiently close to the true value. The results of this approach are presented in figure 11 and they indicate that generally, matrices with fewer interactions require larger numbers of randomizations than matrices with many interactions.

**Figure 11:**
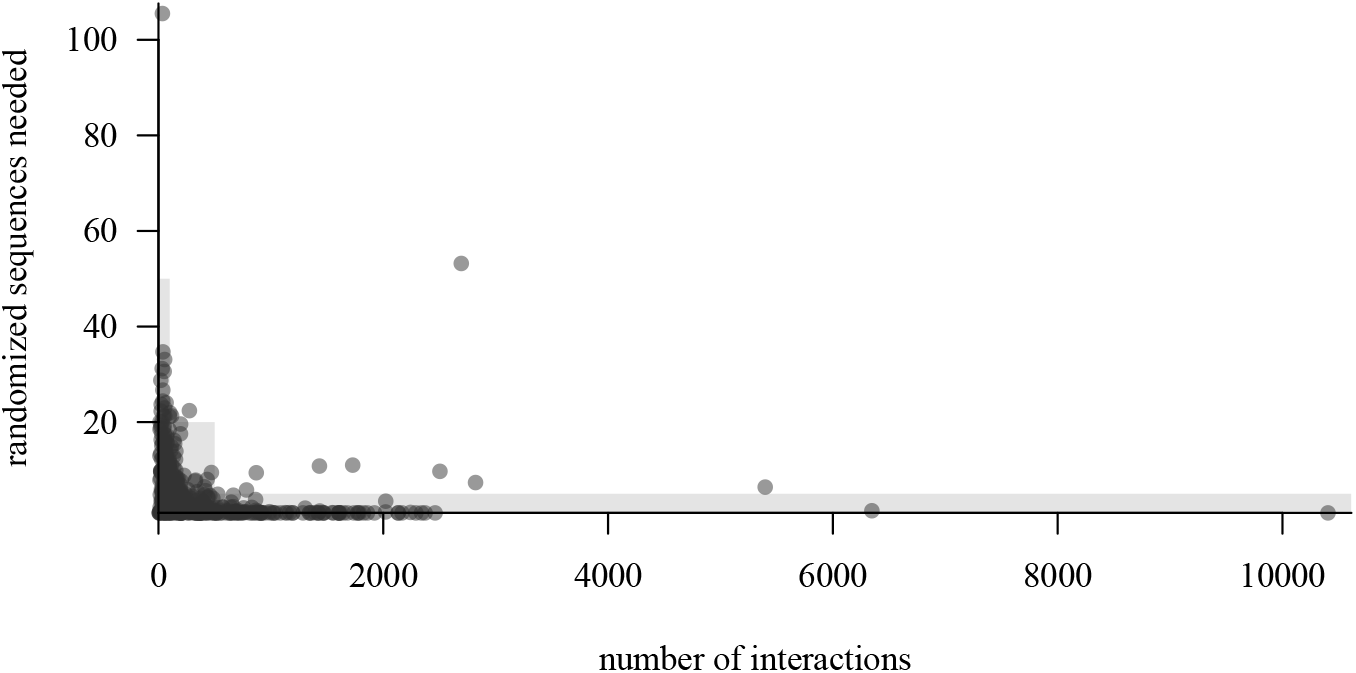
Number of randomized sequences needed to achieve stable results. Matrices with fewer interactions require more randomizations than larger matrices. The shaded areas represent the rule of thumb we used in our evaluations. The majority of matrices achieved stable results when applying these cut-offs.

To be clear, this analysis just serves as a rough guide for setting the number of randomized sequences in our method evaluation. And it is important to remember that whereas our new method provides actual posterior distributions, we had to resort to using point estimates simply for being able to compare our method’s results with the results of the alternative methods. In practice, or if in doubt, it is advisable to set the number of randomizations higher than what we used here and 100 seems a fairly safe value.

## Conclusion

We set out to develop a method that allows unbiased and precise estimation of dominance hierarchy steepness with explicit uncertainty assessment. Our results demonstrate that STEER, steepness based on Bayesian Elo-rating via cumulative winning probabilities, provides such a measure. By also providing the `EloSteepness` R package we make this assessment as user-friendly as possible.

## Appendix

### The problem with classic steepness

**Figure A1:**
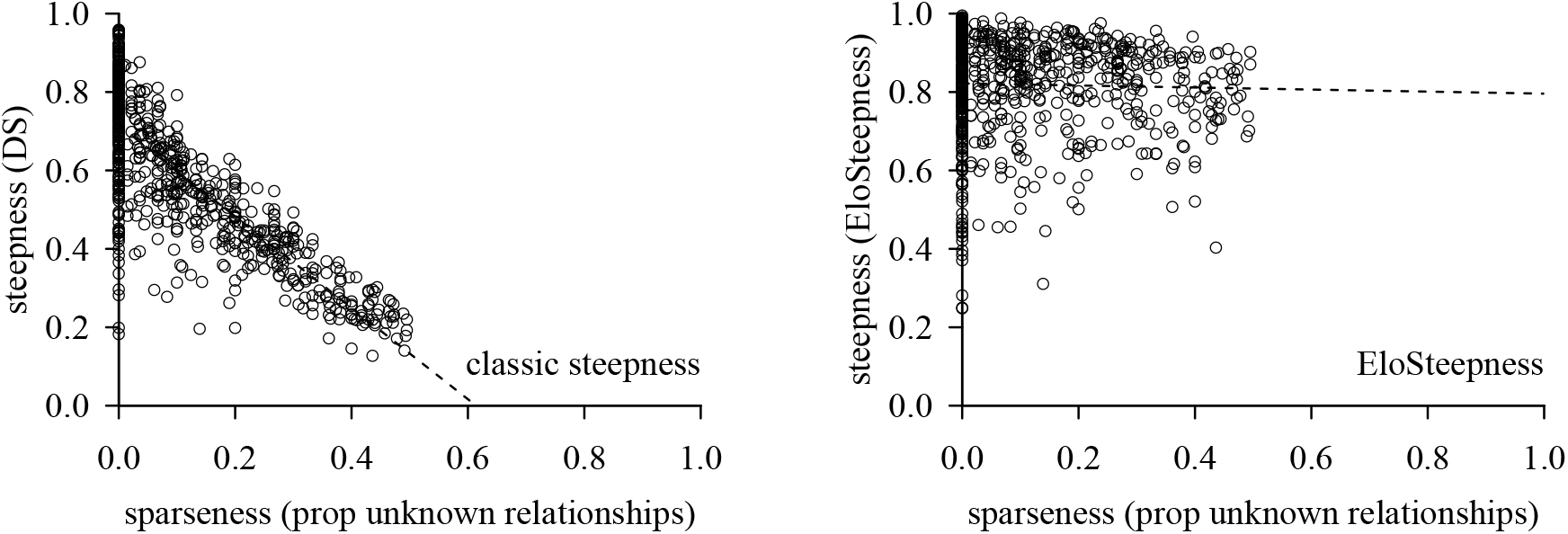
The relationship between data set sparseness and steepness metrics. On the left, classic steepness and its close dependence on unknown relationships. On the right our new STEER metric. Data are from 670 empirical interaction matrices. Dashed lines are fits from simple linear regressions.

### Generating artificial data

First, we evaluated how well our data generation reflects variation in steepness. To this end, we used our 1,000 artificial data sets and selected only those where the initial matrix had less than 5% unknown relationships (494 initial matrices). For these matrices, we correlated the input steepness (i.e., the steepness value we set when the matrix was generated) and the actual observed steepness (based on *P*_*ij*_ David’s scores). See figure A1.

**Figure A2:**
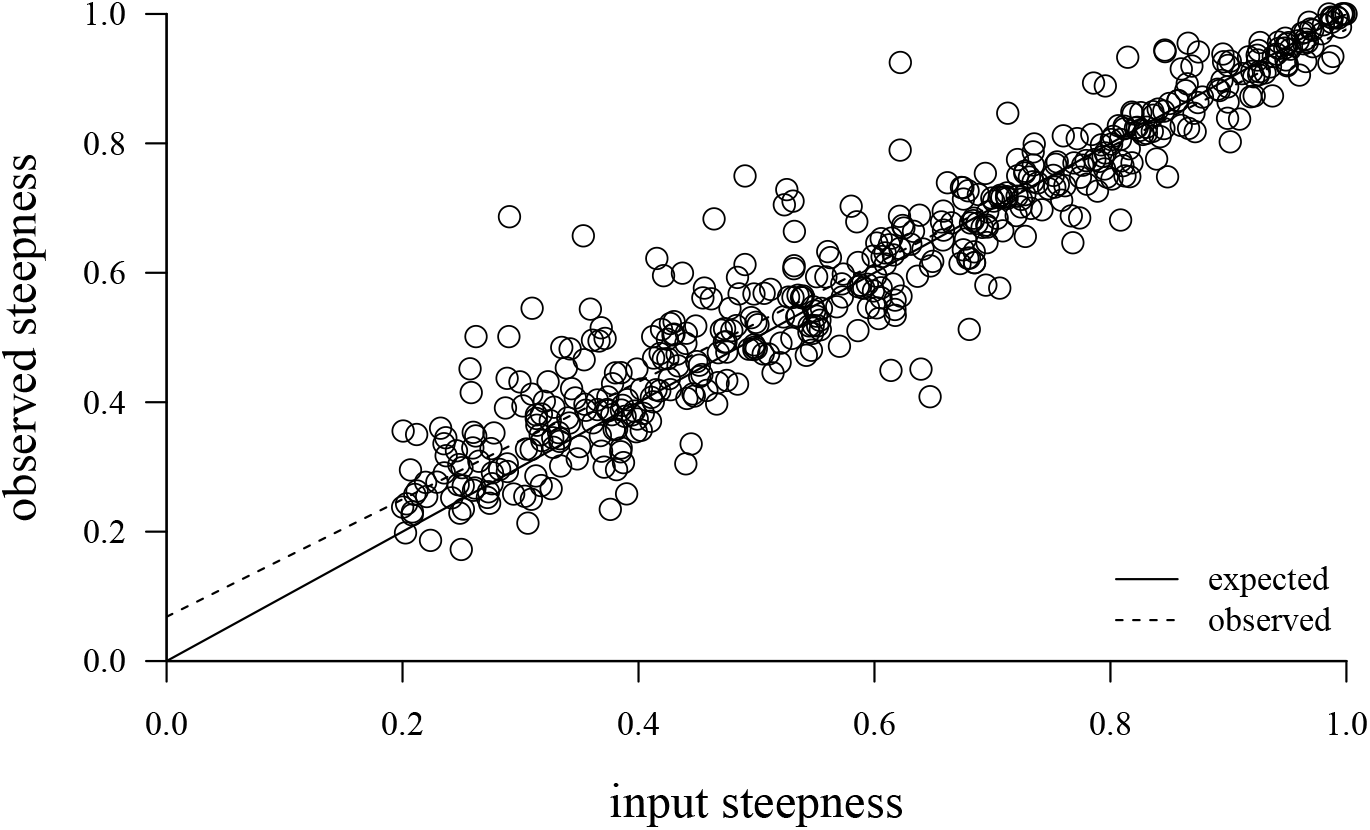
Data generation evaluation. The artificial matrices we generated reflected input steepness fairly well. Each circle reflects one generated matrix (*n* = 494 matrices with no more than 5% unknown relationships).

**Figure A3:**
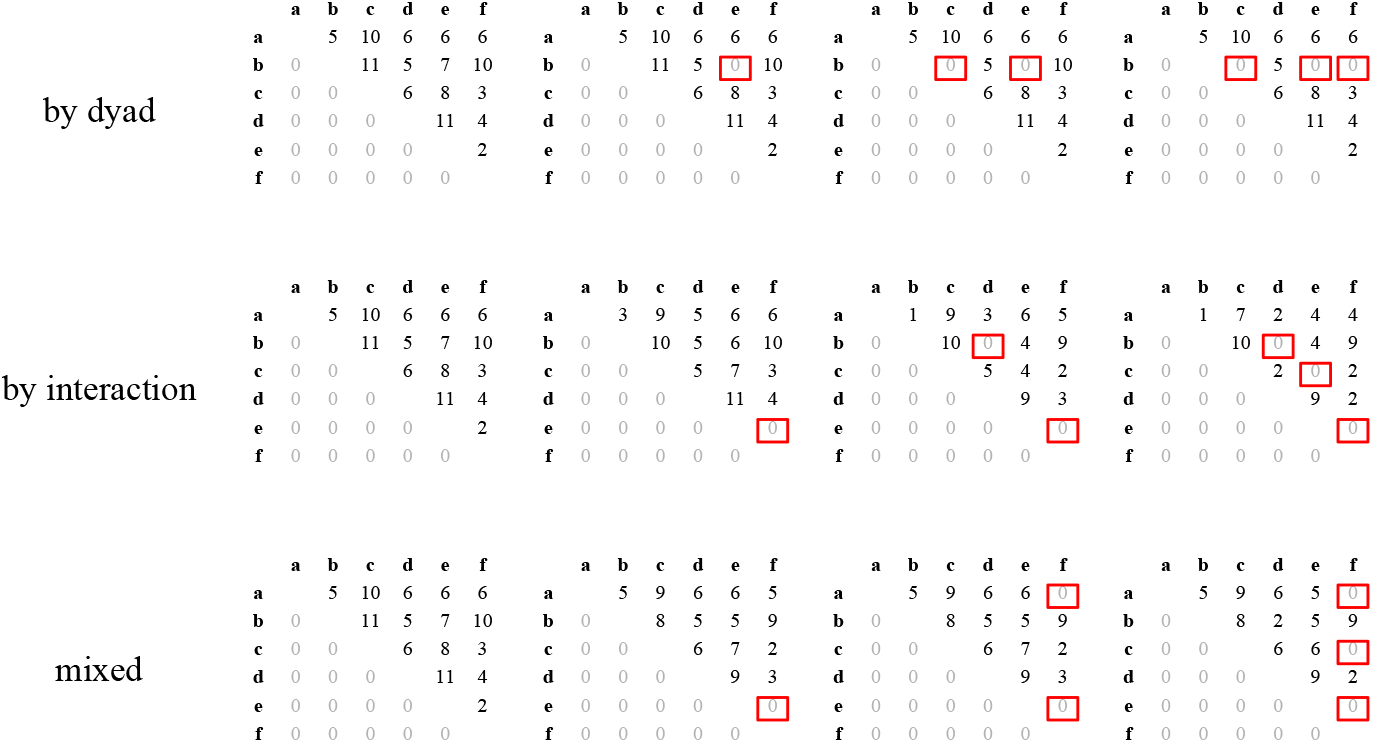
Removal experiment. Starting from the initial matrix (left column), interactions are removed so that the number of unknown relationships increases. The top row shows the approach where entire dyads are set to 0 interactions. The middle row shows how removing interaction by interaction increases the unknown relationships. The bottom row shows a mix between the two. Unknown relationships are highlighted by red boxes in the upper triangles.

### Removal experiments

Setting entire dyads to zero has been done, for example, by Klass & Cords (2011).

### Comparative example revisited

Here we replicate the analysis of Balasubramaniam et al. (2012a) with the data provided in their paper (their table 2), i.e., using point estimates of classic steepness. We reverted to using point estimates because it was not possible to subject all the data sets used in their paper to our new algorithm because the relevant raw data were not available for all data sets.

**Figure A4:**
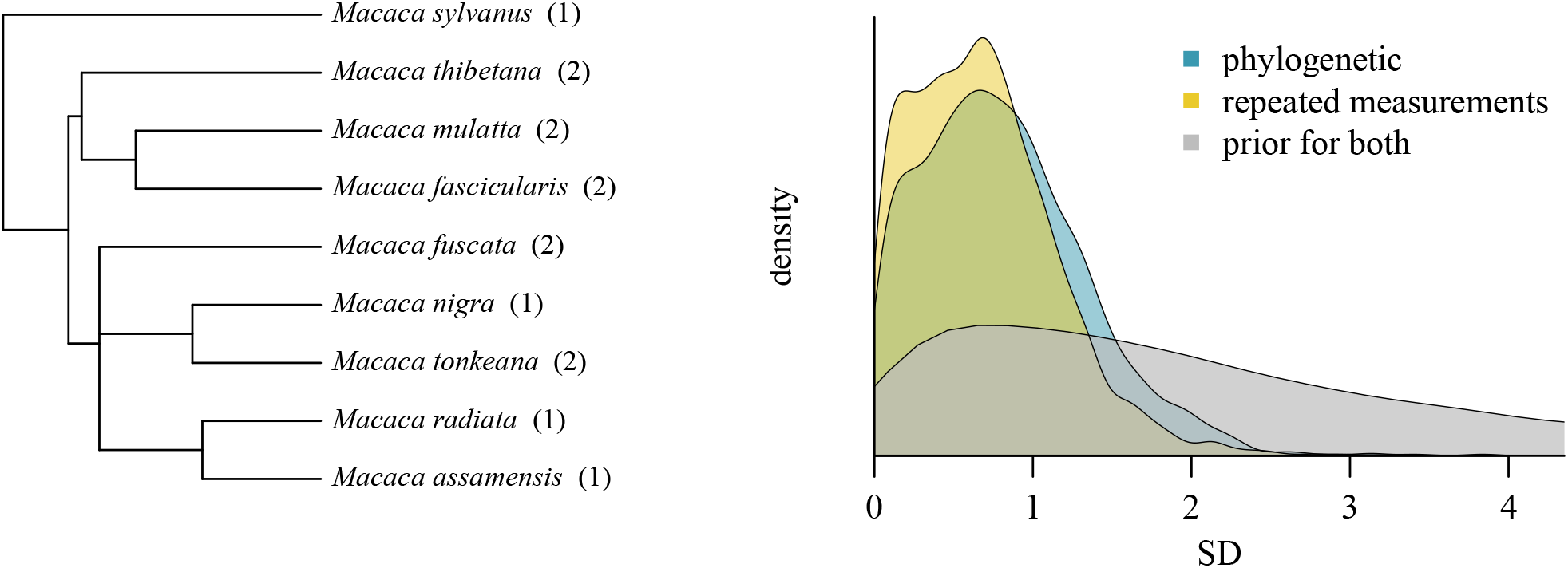
The example from the main text, but fitted to the same data as used by Balasubramaniam et al., i.e., 14 groups of 9 species, using point estimates of classic steepness. In this reanalysis there is somewhat larger phylogenetic variance (mean SD = 0.81) compared to the within species variance (mean SD = 0.68), which is more in line with the conclusions reported in their paper.

### Bayesian David’s scores

Our implementation of David’s scores is coded in `Stan`. Here we model the winning proportion of one individual within a dyad based on the observed outcomes of wins and losses for both dyad members (see figure A5). We model this as a binomial distribution:

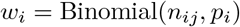

where *w*_*i*_ is the number of wins for an individual within a dyad and *n*_*ij*_ the total number of interactions in the dyad. The winning proportion of the other individual in that dyad is *w*_*j*_ = 1 *− w*_*i*_.

**Figure A5:**
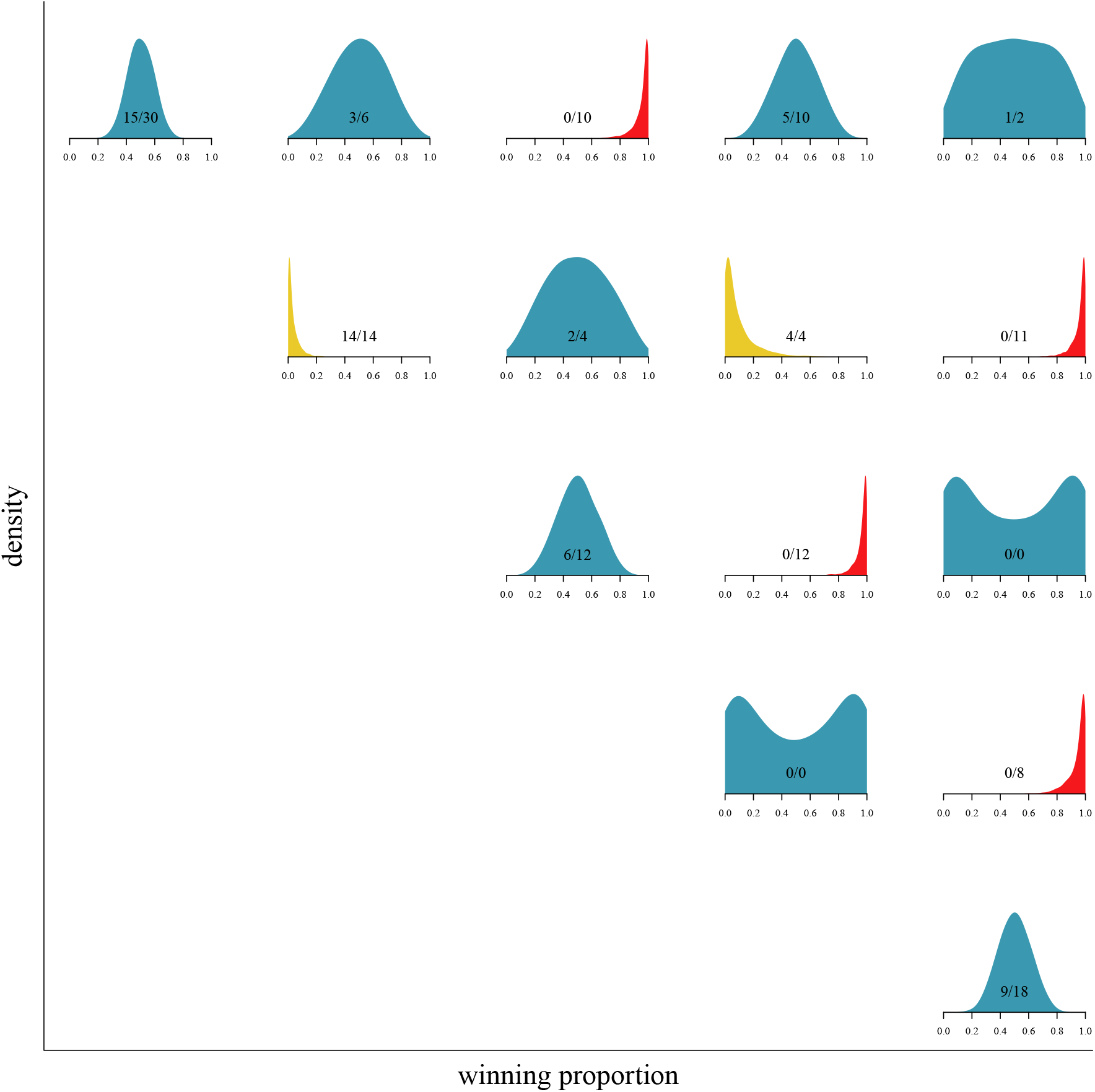
Posteriors of dyadic winning proportions for one member of each dyad in a group of six individuals. Blue distributions indicate individuals in tied or unknown relationships. Red indicates that the depicted individual won all its interactions, and yellow that the individual lost all its interactions. The more interactions were observed (inset text indicates wins/total interactions) the narrower the distributions are.

We put a fairly uninformative, but not flat, prior on *p*, which attributes most weight on winning proportions between 0.05 and 0.95, centered around 0.5 (or more formally: 1/1 + *exp*(−Normal(0, 3)) (see also de Vries et al., 2006) (figure A6). Therefore, if no interactions were observed at all in a dyad, the posterior distribution will reflect the prior and assign winning proportions centered around 0.5. If interactions were observed, the posterior will reflect that and move its density towards the empirical proportion. Importantly, the more interactions were observed the narrower the posterior will become.

**Figure A6:**
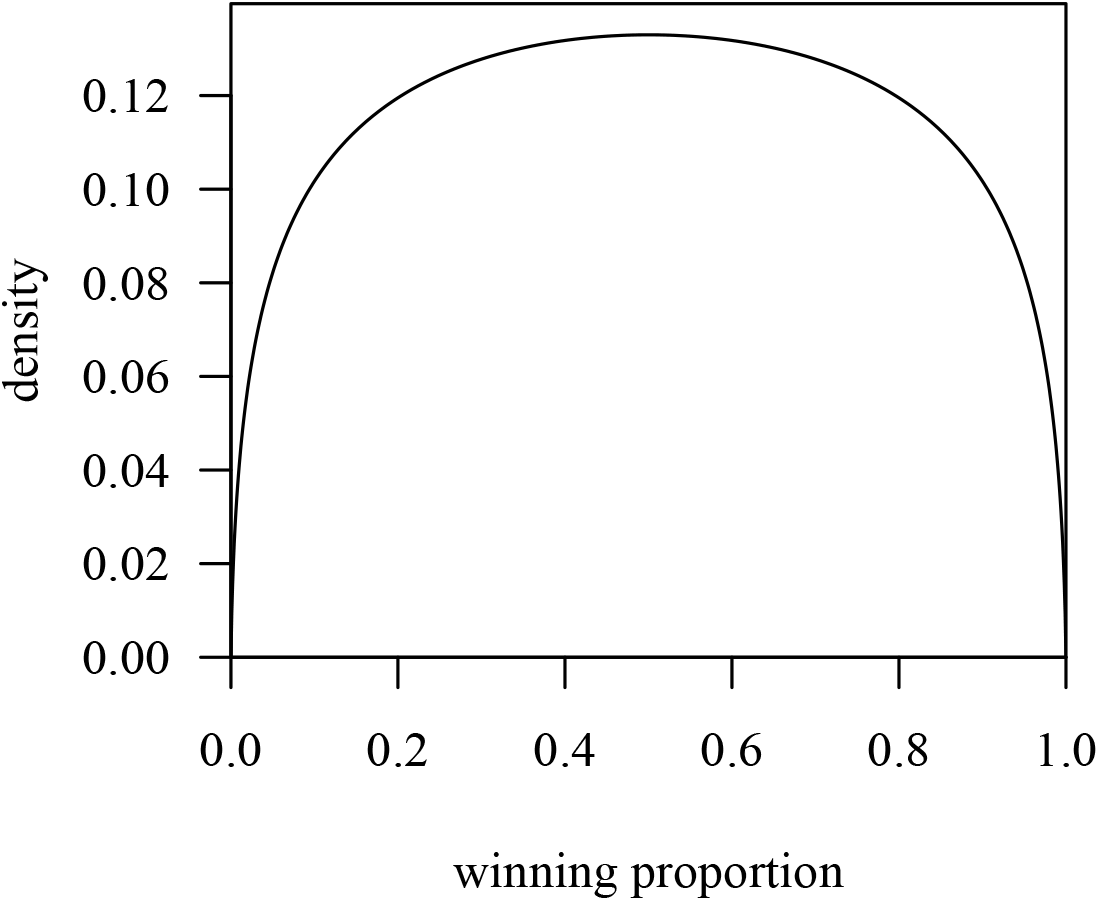
Prior distribution for modelled winning proportions. The prior is fairly diffuse and mostly concentrates its density around a winning proportion of 0.5.

The remaining steps for calculating steepness (summing winning and losing proportions and slope calculation) follow the procedure by de Vries et al. (2006), with the only difference that instead of point estimates we used the full sampling distributions for these calculations, which ultimately also results in a probability distribution for the steepness metric itself.

### Stan implementation

There was one crucial change that we made to the implementation and code provided by Goffe et al. (2018). Instead of estimating the spread of the initial (start) ratings, we kept this parameter fixed. Below in figure A7 we show that the two approaches are in essence indistinguishable. Our modification has the advantage of being computationally faster and resulted substantially less often in divergent transitions.

**Figure A7:**
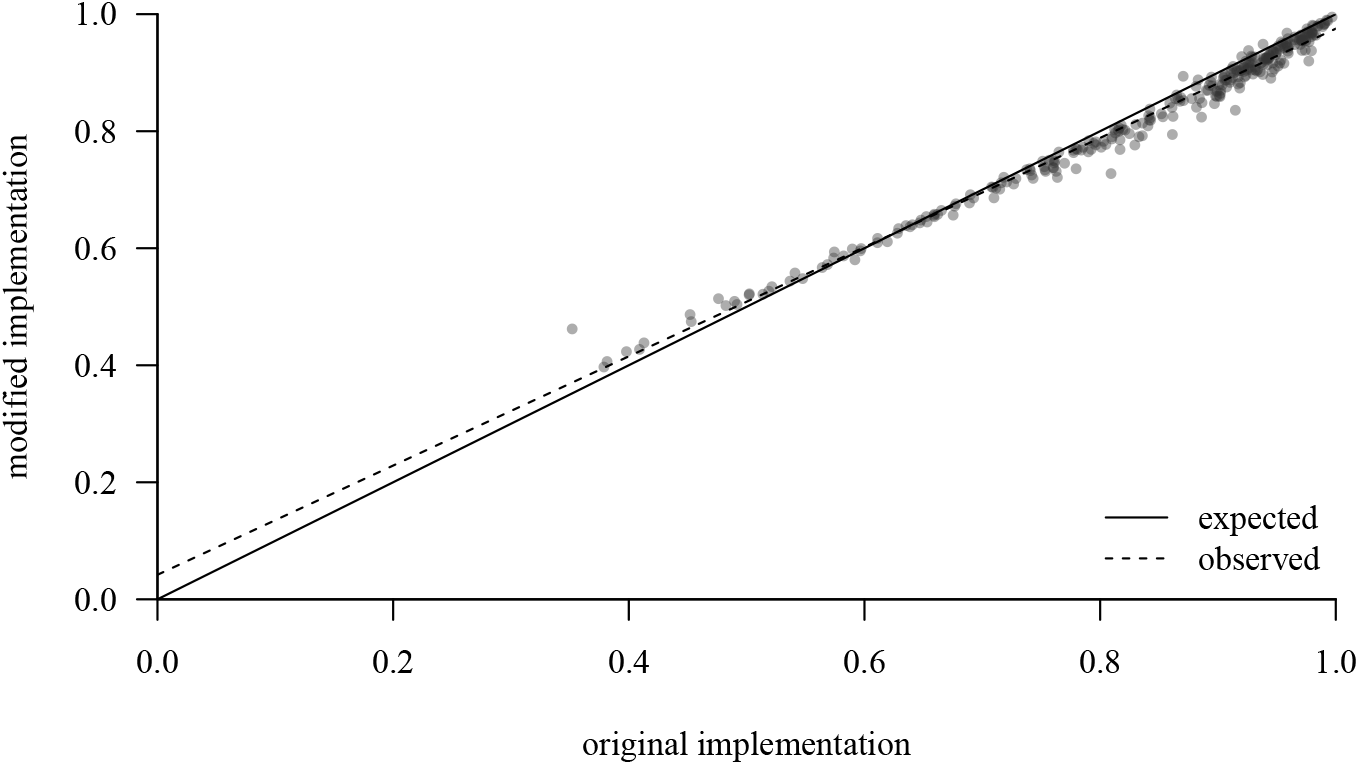
Differences between alternative implementations. Artificial and empirical data are combined here. (total *n* = 300)

## Acknowledgments

We thank Han de Vries and Andrew Robbins for the initial impetus to start this work, and Roger Mundry and Julie Duboscq for numerous discussions. We also thank Han de Vries, Andrew Robbins, David McDonald and Cédric Girard-Buttoz for helpful comments on earlier drafts of this manuscript.

Funding was provided by the German Research Foundation project RTG 2070 Understanding Social Relationships (project number 254142454).

## Footnotes

1 Note that there is a typo in their equation, which we corrected here. Their expression (equation 7) is actually *p*_*BA*_, i.e., the winning probability of *B* against *A*. We corrected this by switching *Elo*_*A*_ and *Elo*_*B*_ in the equation’s denominator.

2 This has direct consequences for the steepness model: in about 50% of MCMC samples *d* has a higher cumulative winning probability than *e* and hence *d*’s ordinal rank in those samples is 4 as opposed to *e*’s rank 5. In the remaining MCMC samples, the ranking of the two is reversed.

